# PREDATION CAPACITY OF SOIL-DWELLING PREDATORY MITES ON TWO GROUND BEETLE IMMATURE STAGES: A BIOLOGICAL CONTROL PERSPECTIVE FOR TWO MAJOR MAIZE PESTS

**DOI:** 10.1101/2019.12.13.862466

**Authors:** Antoine Pasquier, Thibault Andrieux, Paloma Martinez-Rodriguez, Elodie Vercken, Maxime Ferrero

## Abstract

Soil-dwelling predatory mites already proved their efficiency as biocontrol agents against many pests (Carrillo et al. 2015). Western Corn Rootworm (WCR), (*Diabrotica virgifera virgifera)* and Wireworm (WW) (*Agriotes sordidus*) are important pests of various crops (Furlan et al. 2002; Krysan et al., 1986; Ritter and Richter 2013; Wesseler and Fall 2010) whose eggs and first instar larvae also inhabit the first centimeters of soil (Furlan 2004; Vidal et al., 2005). In order to evaluate the potential of predatory mites as biological control agents against WCR and WW, we investigated the predation capacity of *Stratiolaelaps scimitus*, *Gaeolaelaps aculeifer* and *Macrocheles robustulus* on immature stages of these two prey species. First, we observed if one or more species could feed upon eggs and first instar larvae as Prischmann et al. (2011) suggested for WCR. We then explored their predation abilities through time using a survival analysis to identify the best biocontrol agent among the species tested.

Surprisingly, none of the predator species tested identified WCR or WW eggs as preys. However, at least 50% of WCR and WW first instar larvae have been attacked by *G. aculeifer* and *M. robustulus*. *Stratiolaelaps scimitus* showing a slightly lower efficiency (30% of preys attacked). The survival analysis confirmed this trend with slower predation dynamics for *S. scimitus*.

These results show a potential of soil-dwelling predatory mites as biocontrol agents against WCR and WW. Furthermore, targeting the neonate stage instead of the egg stage in pest management strategies seems necessary for maximizing efficiency.

## INTRODUCTION

Western corn rootworm (WCR) (*Diabrotica virgifera virgifera*) (LeConte, 1868) (Coleoptera: *Chrysomelidae*) and Wireworm (WW) (*Agriotes sordidus*) (Illiger, 1807) are two corn (*Zea mays* L.) pests present in Europe and North America (Kriticos et al. 2012; Platia 1994). Krysan et al. (1986) estimated WCR economic impact in North America over 1 billion USD in 1986, including yield losses and pest management. In Europe, Wesseler and Fall (2010) valued the economic impact in case of no control of the pest at 700 million euros per year. *Agriotes sordidus* economic impact is not well-estimated but is considered as one of the most harmful *Agriotes* species from an agronomic point of view (Furlan 2004). Even if adults can feed on silk, leaves and pollen (Elliott, Gustin, et Hanson 1990; Lehmhus et Niepold 2013), main losses are caused by WCR and WW larval stages. They feed on underground parts of the plant (Furlan 2004; Vidal et al., 2005), reducing crops productivity, and increasing risk of lodging for WCR (Furlan 2004; Levine et al. 2002). WCR is more specific than WW with corn being the main crop impacted, even if one population became recently a pest for soybean (Spencer et al. 2009; Curzi et al. 2012; Gray et al. 2009). *Agriotes sordidus* is considered a generalist pest, causing economic damages on many vegetable and arable crops, including potato, sugar beet, corn, tomato, onion, watermelon and melon (Burgio et al. 2012).

Some pest management tools already exist to control WCR populations: pesticides, Genetically Modified Organisms (GMOs) producing the *Bacillus thuringiensis* (Beliner, 1915) toxin, or crop rotation. In all cases, at least one WCR population evolved and became resistant to the pest control methods. Pesticides (Pereira et al. 2017), GMOs (Gassmann et al. 2011) and soybean/corn crop rotation became obsolete pest management methods in the areas concerned. Barsics et al. (2013) detailed the management pest methods available on *Agriotes spp.*. They highlighted the need of a better Integrated Pest Management (IPM) as recommended by the European and Mediterranean Plant Protection Organisation (EPPO) in 2014 and in the recommendation 2014/63/EU for WCR. Moreover, the commission implementing regulation 2013/485/EU is restricting the use of neonicotinoids. These pesticides being widely use against the two pests previously mentioned makes their control even more challenging.

In this context, Prischmann et al. (2011) opened a new perspective to control WCR by showing a predation activity of soil-dwelling predatory mites species (*Gaeolaelaps aculeifer*) (Canestrini, 1883) on WCR eggs and first instar larvae. Indeed, soil-dwelling predatory mites that inhabit the first centimeters of the soil (Moreira et de Moraes 2015) have been widely used to control pests such as nematodes, thrips or flies (Carrillo et al. 2015). Three species are already commercialized for their interest in biological control: *Gaeolaelaps aculeifer*, *Stratiolaelaps scimitus* (Womersley) and *Macrocheles robustulus* (Berlese).

The purpose of this study is to experiment new potential targets for these generalist soil-dwelling predators by testing their predation activity on WCR and WW immature stages.

## MATERIALS AND METHODS

### PREDATORY MITES

The three species used in this experiment were stored in climatic chambers at 25°C +/− 0,5°C and 70% +/− 10 RH% with a constant obscurity. A mix of *Aleuroglyphus ovatus* stages were used as food and extra water were provided three times a week in a 100 mm × 94 mm bugdorm-5002 with 30μm nylon screen port sold by Bugdorm©.

#### Macrocheles robustulus

Koppert Biological systems provided *Macrocheles robustulus*. Their product is called Macro-mite©. We maintained them on vermiculite during 2 months with a mix of *A. ovatus* stages.

#### Gaeolaelaps aculeifer

*Gaeolaelaps aculeifer* is produced by EWH Bioproduction, Denmark. The population was maintained during 8 months on a substrate made of 1/3 third blond sphagnum peat and 2/3 of fine vermiculite and fed with a mix of *A. ovatus* stages.

#### Stratiolaelaps scimitus

*Stratiolaelaps scimitus* individuals used in this experiment are produced by Bioline AgroSciences. The product is called Hypoline©. This population has been maintained on blond sphagnum peat and fed with a mix of *A. ovatus* stages for 2 years.

### PREYS

We experimented eggs and first instar larvae for both *Diabrotica virgifera virgifera* and *Agriotes sordidus* as potential preys. We also added *Aleuroglyphus ovatus* eggs as a positive control of predation activity since astigmatid mites are known to be a suitable food source for those species (Rueda-Ramirez et al. 2018).

#### *Diabrotica virgifera virgifera* eggs

WCR diapausing eggs were provided by the Centre of Agriculture and Bioscience International (CABI), Hungary. They were stored at 7°C +/− 0,5°C below their temperature of development (Meinke et al. 2009). We sieved the eggs them from their substrate and selected only turgescent eggs to offer them to the predatory mites.

#### *Diabrotica virgifera virgifera* first instar larvae

We placed WCR eggs on plaster of Paris in a climatic chamber at 25°C +/− 0,5°C and 70% +/− 10 RH%. We added water twice a week to keep the plaster of Paris moist. We checked daily if eggs hatched and introduced the first instar larvae in the predation device.

#### *Agriotes sordidus* eggs

Arvalis provided *Agriotes sordidus* eggs and first instar larvae by sending us couple of adults ready to lay eggs in Petri dishes filled with a sample of soil where they have been collected. Both eggs and first instar larvae have been extracted from this dirt.

#### *Aleuroglyphus ovatus* eggs

*A. ovatus* eggs are produced by Bioline AgroSciences. Eggs were sterilized before presentation to the predatory mites.

### PREDATION DEVICE

Predation tests have been inspired from El Adouzi, Bonato, et Roy 2017; Lovis et al. 2011 and Nordenfors et Hoglund 2000 protocols by isolating each mite individually. However, we chose to carry out the predation tests in 2 mL Eppendorf tubes containing each 1 mL of dried plaster of Paris to maintain a high percentage of humidity necessary to soil-dwelling predatory mites survival (El Adouzi, Bonato, et Roy 2017). Adult mites of both sexes were individually isolated and starved during 7 days in the tubes before the predation tests. In total, 240 predatory mites have been isolated with 1/3 of each species to present them to 4 different types of preys. Twenty predation tests were made by prey/predator couple.

During the 7-days period of starvation we added 100μL of water every 3 days to maintain a suitable relative humidity necessary for soil-dwelling predatory mites survival. We also drilled the top of the tube and covered it with a 106 μm mesh width nylon tissue. This size of mesh allowed for water and gas exchange while preventing mites from leaving the tube. These tubes were stored in a climatic chamber at 25°C +/− 0,5°C with 70% +/− 10% RH. All three species were active after this period of storage and starvation.

We introduced 20 times one prey in a tube containing a predatory mite and observed predation activity during maximum 10 minutes or less if predation happens before that timing. We observed each mite feeding or non-feeding activity through the tube with a binocular. We used an indirect source of light, controlled at 100 lux (measured with the Digital Illuminance meter TES 1335), to minimize natural behavior disruption of these lucifugous species. During each assay, the timing and number of contacts between the predator and the prey before predation were noted. We considered predation activity when mites impaled the prey with their chelicerae. We chose to observe predation on a short duration because some of the preys could be impacted by plaster of Paris abrasive texture if it dries up.

### STATISTICAL ANALYSES

All analyses were done using R (R Core Team 2018). Predation rate was analyzed with a post-hoc analysis on a Fisher’s exact test, based on **rcompanion** package in R (Mangiafico 2018). Considering that multiple pair wise test can induce false-positive results, we used a Bonferroni correction to minimize this error to analyze and rank the *p*-values.

Time to predation on WCR and WW first instar larvae was analyzed with a Log-rank test pairwise comparison (**survival** package, Therneau 2015).

## RESULTS

### PREDATION RATE

The three species of predatory mites tested on *A. ovatus* eggs showed a high predation rate with 100% +/−0%, 90% +/− 31% and 80% +/− 41% of predation for *M. robustulus*, *G. aculeifer* and *S. scimitus,* respectively. *M. robustulus* and *G. aculeifer* seem to show a higher predation rate on WCR and WW first instar larvae compare to *S. scimitus* (Figure 1) but it is not statistically supported (*G. aculeifer/ S. scimitus* on WCR first instar larvae *p*-value=0,207; *M. robustulus / S. scimitus* on WCR first instar larvae *p*-value=0,536; *G. aculeifer/ S. scimitus* on WW first instar larvae *p*-value=0,536*; M. robustulus / S. scimitus* on WW first instar larvae *p*-value=0,536). WCR and WW eggs were never attacked by *S. scimitus* and *M. robustulus*, and in only 10% +/− 31% of the assays by *G. aculeifer* (Figure 1) on WCR and none for WW.

**Fig 1 :**
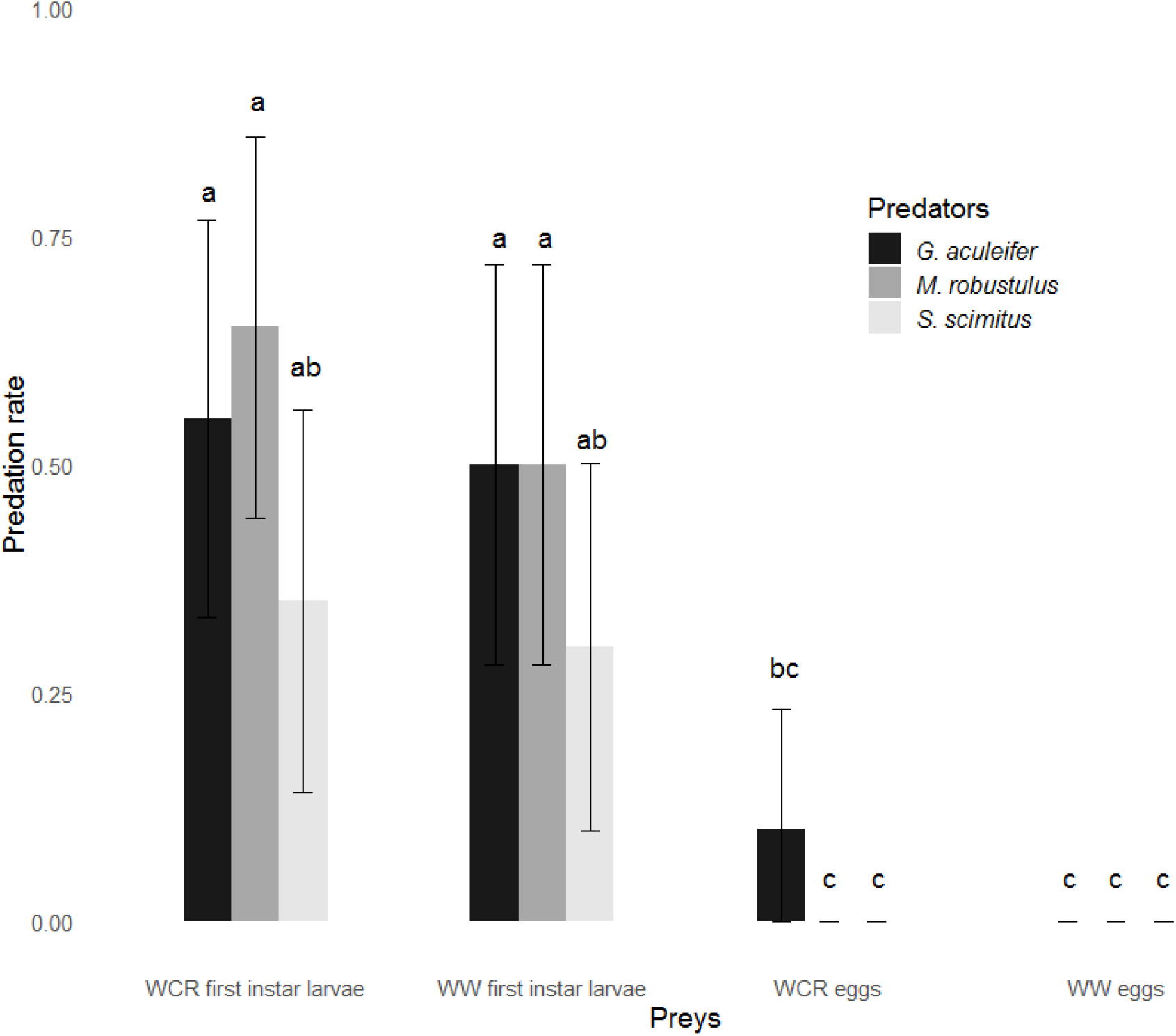
Predation rate of *G. aculeifer*, *S. scimitus* and *M. robustulus* on WCR and WW eggs and first instar larvae during the 10-minutes predation assays (20 tests per prey/predator couple). We used a Fischer’s exact test with Bonferroni correction to analyze the data. Significant groups (with a *p*-value<0,05) are represented by letters on top of each bar. The error bars represent the 95% confidence interval for the predation rate.

### SURVIVAL ANALYSIS

The log-rank pairwise comparison confirm the trend that *S. scimitus* is less efficient to find and attack WCR first instar larvae compared to *M. robustulus* (*p*-value = 0,073) and *G. aculeifer* (*p*-value = 0,169, Fig.2A). However, *S. scimitus, M. robustulus* and *G. aculeifer* predation efficiency to feed upon the WW first instar larvae is statistically the same (Fig.2B, *M. robustulus/G. aculeifer*: *p*-value= 0.51, S. *scimitus/G. aculeifer*: *p*-value= 0.39, *M. robustulus/S. scimitus*: *p*-value= 0.25). Out of the statistical analysis, we can observe that most of the predation events occurs in the first 200s of observation except for *G. aculeifer* on WW where it happens all along the 10 min of observation.

**Fig 2.**
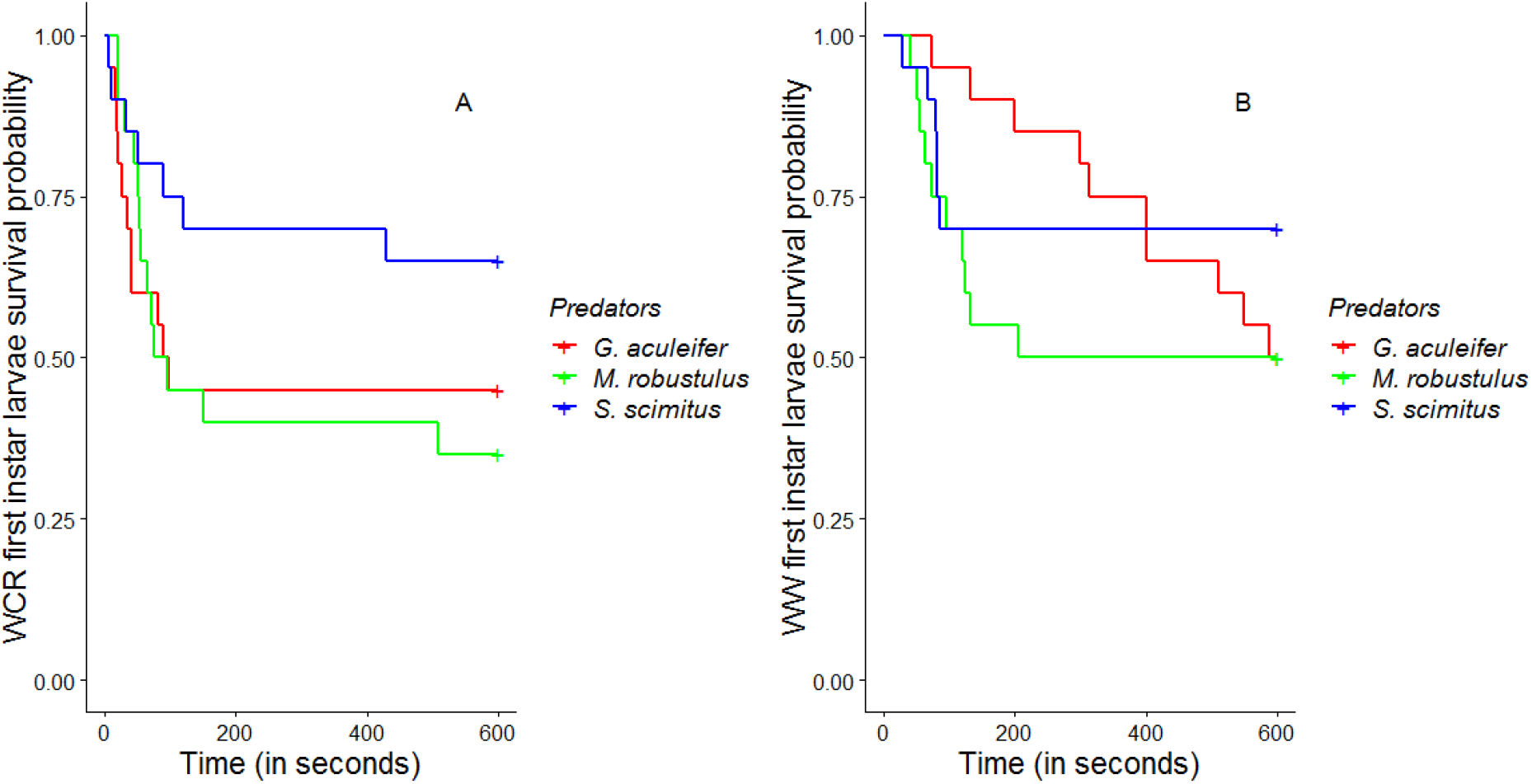
First instar WCR and WW larvae survival probability in presence of soil-dwelling predatory mites species through time (for n=20 predation test) represented by survival curve. No significant differences have been identified using a significant threshold with a *p*-value>0.05.

## DISCUSSION

In this study, we wanted to investigate the biological control potential of soil-dwelling predatory mites on WCR and WW. Our experiment demonstrated that three species commonly used in biological control can effectively prey upon one of the first stage of development of WCR or WW, and that two of them (*M. robustulus* and *G. aculeifer)* have maximum efficacy to prey upon these ground beetle larvae in our experimental conditions.

First, we controlled that our conditions of experimentation were compatible with the specific ecology of the predatory mites tested. Indeed, soil-dwelling predatory mites are very sensitive to humidity (El Adouzi, Bonato, et Roy 2017) and our will to have a period of 7 days of starvation required to maintain a high percentage of humidity in the predation device to keep them alive. Our capacity to keep in total 300 predatory mites active for 7 days using this predation device shows its ability to maintain this ecological requirement. Moreover, soil-dwelling predatory mites are lucifugous and our wish to record timing of predation required to have at least a source of light. In order to verify if our light conditions inhibit the predation behavior of predatory mites, we observed predation activity on a known prey routinely consumed by all three species (Rueda-Ramirez et al. 2018). Our results confirm that the experimental conditions do not inhibit predation since we observed 80 +/− 41%, 90 +/− 31% and 100 +/− 0% of predation within 10 minutes of observation for *S. scimitus*, *G. aculeifer* and *M. robustulus*, respectively.

Consecutively to the control of the validity of our experimental design, we can affirm that all three species of the predatory mites tested are unable to identify eggs of WCR and WW as preys. These results are in contradiction with Prischmann et al. (2011) observations on the use of *G. aculeifer* on WCR which show positive results. We observed once an attack from a *G. aculeifer* on a damaged egg that we took off from the data set. We also carried out a complementary experiment to verify whether damaged eggs could be attacked and consumed. We found that *G. aculeifer* is very efficient to attack WCR damaged eggs with 85% +/− 37% of predation following the same methods presented in this study (supplementary materials). We hypothesize that quality of the eggs and/or the duration of experimentation (24 hours for Prischmann et al. (2011)) could be the origin of the difference in our results.

All three predator species tested have been able to feed upon WCR and WW first instar larvae. These results confirm Prischmann et al. (2011) conclusion on this development stage for *G. aculeifer*. However, the survival analysis showed a slower predation activity for *S. scimitus* on WCR leading us to consider *G. aculeifer* or *M. robustulus* instead of *S. scimitus* for future studies on this pest.

These results highlight a potential interest of predatory mites for biological control against ground beetles. However, more work remains to be done in conditions closer to *in-situ* environment to take into accounts more parameters of the complexity of the predator prey interactions. Indeed, we highlighted the capacity of predatory mites to identify and attack a special kind of prey. However, Abrams (2000) enhances the effect of many parameters such as environment effect or population density which could impact the actual mites predation efficiency on ground beetle larvae. In the perspective of using soil-dwelling predatory mites as biocontrol agents of WCR or WW, our results also emphasize the need to focus on a specific developmental timing to have a chance to control the damages on plant. Indeed, eggs are not consumed by the predatory mites and the WCR first instar larvae can establish near the roots as soon as they find one (Spencer et al. 2009). Moreover, Lundgren et al. (2009) suggest that WCR older stage are able to protect themselves against predators while attacked thanks to their hemolymph properties. Indeed, their hemolymph, once in predators’ mouthparts, will coagulate and then stop or slow down the predation activity. Other defensive mechanisms are present for WW, *e.g.* older larvae stage of development have a sclerified cuticle that likely protect them against mites attack (Chaton et al. 2008).

The conditions just listed are restrictive for the use of biocontrol agents. If the favorable timing to control the pest is restricted to the few days between the eggs hatching and the establishment of larvae near the root (Macdonald et Ellis 1990), the agent should have maximum efficiency on the sensitive stage that is first instar larvae. In our experimental conditions, while all three species seem to be candidates to control WW, we observed a lower potential for biological control from *S. scimitus* on WCR compared to *M. robustulus* and *G. aculeifer.* However, although we have demonstrated that soil-dwelling predatory mites have the potential to attack and predate WCR and WW first instar larvae, it is now necessary to confirm this result *in situ* or in semi-controlled conditions experimentations to check whether predatory mites will effectively be able to find and control the first instar larvae of these pests in the soil.

## Supporting information

Predation tests on WCR broken eggs

## ACKNOWLEDGEMENTS

We are thankful to the French Interprofessional Organisation for Seeds and Plants (GNIS) which enabled this scientific work by funding the Mites Against Diabrotica (M.A.D.) project. We also acknowledge all the partners in this project: Axereal Serbia, French National Institute for Agricultural Research (INRA) and with a more specific attention Arvalis, which provided *Agriotes sordidus* eggs and larvae, essential for this study.

## REFERENCES

2013/485/EU. 2013. « COMMISSION IMPLEMENTING REGULATION (EU) No 485/2013 of 24 May 2013 Amending Implementing Regulation (EU) No 540/2011, as Regards the Conditions of Approval of the Active Substances Clothianidin, Thiamethoxam and Imidacloprid, and Prohibiting the Use and Sale of Seeds Treated with Plant Protection Products Containing Those Active Substances ». 2013. https://eur-lex.europa.eu/eli/reg_impl/2013/485/oj.

2014/63/EU. 2014. « Recommandation n° 2014/63/UE de la Commission du 06/02/14 relative à des mesures de lutte contre *Diabrotica virgifera virgifera* Le Conte dans les zones de l’Union où sa présence est confirmée | AIDA ». 2014. https://aida.ineris.fr/consultation_document/32035.

Abrams, P. A. 2000. « The Evolution of Predator-Prey Interactions: Theory and Evidence ». Annual Review of Ecology and Systematics 31 (1): 79–105. https://doi.org/10.1146/annurev.ecolsys.31.1.79.

Barsics, F., E. Haubruge, et F. Verheggen. 2013. « Wireworms’ Management: An Overview of the Existing Methods, with Particular Regards to *Agriotes Spp. (Coleoptera: Elateridae)* ». Insects 4 (1): 117–52 https://doi.org/10.3390/insects4010117.

Burgio, G., G. Ragaglini, R., Petacchi, R. Ferrari, M. Pozzati, et L. Furlan. 2012. « Optimization of *Agriotes sordidus* monitoring in northern Italy rural landscape, using a spatial approach ». Bulletin of Insectology 65 (1): 123–131.

Carrillo, D., G. José de Moraes, et J. E. Peña, éd. 2015. Prospects for Biological Control of Plant Feeding Mites and Other Harmful Organisms. Progress in Biological Control 19. Cham: Springer.

Chaton, P. F., G. Lemperiere, M. Tissut, et P. Ravanel. 2008. « Biological traits and feeding capacity of *Agriotes* larvae *(Coleoptera: Elateridae)* : A trial of seed coating to control larval populations with the insecticide fipronil ». Pesticide biochemistry and physiology 90 (2): 97–105.

Curzi, M. J., J. A. Zavala, J. L. Spencer, et M. J. Seufferheld. 2012. « Abnormally High Digestive Enzyme Activity and Gene Expression Explain the Contemporary Evolution of a *Diabrotica* Biotype Able to Feed on Soybeans ». Ecology and Evolution 2 (8): 2005–17. https://doi.org/10.1002/ece3.331.

E. Pereira, A., D., Souza, N. S., Zukoff, L. J., Meinke, et B. D., Siegfried,. 2017. « Cross-Resistance and Synergism Bioassays Suggest Multiple Mechanisms of Pyrethroid Resistance in Western Corn Rootworm Populations ». Édité par Raul Narciso Carvalho Guedes. PLOS ONE 12 (6): e0179311. https://doi.org/10.1371/journal.pone.0179311.

El Adouzi, M., O., Bonato, et L., Roy, 2017. « Detecting Pyrethroid Resistance in Predatory Mites Inhabiting Soil and Litter: An *in Vitro* Test: Tarsal Contact Method for Assessing Pesticide Resistance in Mesostigmata ». Pest Management Science 73 (6): 1258–66. https://doi.org/10.1002/ps.4454.

Elliott, N. C., R. D. Gustin, et S. L. Hanson. 1990. « Influence of adult diet on the reproductive biology and survival of the western corn rootworm, *Diabrotica virgifera virgifera* ». Entomologia experimentalis et applicata 56 (1): 15–21.

Furlan, L. 2004. « The Biology of *Agriotes Sordidus* Illiger (*Col., Elateridae*) ». Journal of Applied Entomology 128 (9–10): 696–706. https://doi.org/10.1111/j.1439-0418.2004.00914.x.

Furlan, L., M. Toth, W. E. Parker, M., Ivezić, S. Pančić, M., Brmež, R., Dobrinčić, et al. 2002. « The Efficacy of the New *Agriotes* Sex Pheromone Traps in Detecting Wireworm Population Levels in Different European Countries ». In. https://bib.irb.hr/prikazi-rad?rad=134098.

Gassmann, A. J., J. L., Petzold-Maxwell, R. S., Keweshan, et M. W., Dunbar. 2011. « Field-Evolved Resistance to Bt Maize by Western Corn Rootworm ». Édité par Peter Meyer. PLoS ONE 6 (7): e22629. https://doi.org/10.1371/journal.pone.0022629.

Gray, M. E., T. W., Sappington, N. J., Miller, J., Moeser, et M. O., Bohn. 2009. « Adaptation and Invasiveness of Western Corn Rootworm: Intensifying Research on a Worsening Pest ». Annual Review of Entomology 54 (1): 303–21. https://doi.org/10.1146/annurev.ento.54.110807.090434.

Kriticos, D. J., P. Reynaud, R. H. A. Baker, et D. Eyre. 2012. « Estimating the Global Area of Potential Establishment for the Western Corn Rootworm (*Diabrotica Virgifera Virgifera*) under Rain-Fed and Irrigated Agriculture*: The Potential Distribution of Western Corn Rootworm ». EPPO Bulletin 42 (1): 56–64. https://doi.org/10.1111/j.1365-2338.2012.02540.x.

Krysan, J. L., T. A., Miller, et J. F. Andersen. 1986. Methods for the study of pest Diabrotica. Springer-Verlag.

Lehmhus, J., et F., Niepold. 2013. « Neue Funde des Schnellkäfers *Agriotes sordidus* (Illiger, 1807) mit einem Überblick über seine aktuelle Verbreitung in Deutschland ». Journal für Kulturpflanzen 65(8) 2013, août, 1,15 MB, 309–14. https://doi.org/10.5073/jfk.2013.08.02.

Levine, E., J. L. Spencer, S. A. Isard, D. W. Onstad, et M. E. Gray. 2002. « Adaptation of the Western Corn Rootworm to Crop Rotation: Evolution of a New Strain in Response to a Management Practice. » American Entomologist 48 (2): 94–117.

Lundgren, J. G., T., Haye, S., Toepfer, et U., Kuhlmann. 2009. « A Multifaceted Hemolymph Defense against Predation in *Diabrotica Virgifera Virgifera* Larvae ». Biocontrol Science and Technology 19 (8): 871–80. https://doi.org/10.1080/09583150903168549.

Macdonald, P. J., et C. R. Ellis. 1990. « Survival time of unfed, first-instar western corn rootworm (Coleoptera: *Chrysomelidae*) and the effects of soil type, moisture, and compaction on their mobility in soil ». Environmental Entomology 19 (3): 666–671.

Mangiafico, S., 2018. rcompanion: Functions to Support Extension Education Program Evaluation. (version R package 2.0.3.). https://CRAN.R-project.org/package=rcompanion.

Meinke, L J., T. W., Sappington, D. W., Onstad, T., Guillemaud, N. J., Miller, J., Komáromi, N., Levay, Lorenzo Furlan, József Kiss, et Ferenc Toth. 2009. « Western corn rootworm (*Diabrotica virgifera virgifera* LeConte) population dynamics ». Agricultural and Forest Entomology 11 (1): 29–46.

Moreira, G. F., et G., José de Moraes. 2015. « The Potential of Free-Living Laelapid Mites (*Mesostigmata: Laelapidae*) as Biological Control Agents ». In Prospects for Biological Control of Plant Feeding Mites and Other Harmful Organisms, édité par Daniel Carrillo, Gilberto José de Moraes, et Jorge E. Peña, 77–102. Cham: Springer International Publishing. https://doi.org/10.1007/978-3-319-15042-0_3.

Platia, G. 1994. « Fauna d’Italia, *Coleoptera Elateridae* ». Edizioni Calderini.

Prischmann, D. A., E. M., Knutson, K. E., Dashiell, et J. G., Lundgren. 2011. « Generalist-Feeding Subterranean Mites as Potential Biological Control Agents of Immature Corn Rootworms ». Experimental and Applied Acarology 55 (3): 233–48. https://doi.org/10.1007/s10493-011-9468-y.

R Core Team. 2018. « R: A language and environment for statistical computing. R Foundation for Statistical Computing », 2018. URL https://www.R-project.org/.

Ritter, C., et E., Richter. 2013. « Control Methods and Monitoring of *Agriotes* Wireworms *(Coleoptera: Elateridae)* ». Journal of Plant Diseases and Protection 120 (1): 4–15. https://doi.org/10.1007/BF03356448.

Rueda-Ramirez, D. M., D., M Rios-Malaver, A., Varela-Ramirez, et G. J., de Moraes. 2018. « Colombian population of the mite *Gaeolaelaps aculeifer* as a predator of the thrips *Frankliniella occidentalis* and the possible use of an astigmatid mite as its factitious prey ». Systematic and Applied Acarology 23 (12): 2359. https://doi.org/10.11158/saa.23.12.8.

Spencer, J. L.,, B. E., Hibbard, J., Moeser, et D. W., Onstad. 2009. « Behaviour and Ecology of the Western Corn Rootworm (*Diabrotica Virgifera Virgifera* LeConte) ». Agricultural and Forest Entomology 11 (1): 9–27. https://doi.org/10.1111/j.1461-9563.2008.00399.x.

Therneau, T. 2015. A Package for Survival Analysis in S (version 2.38). <URL: https://CRAN.R-project.org/package=survival>.

Vidal, S., U., Kuhlmann, et C. R., Edwards, éd. 2005. Western corn rootworm: ecology and management. Wallingford, Oxfordshire ; Cambridge, MA: CABI Pub.

Wesseler, J., et E. H. Fall. 2010. « Potential Damage Costs of *Diabrotica Virgifera Virgifera* Infestation in Europe - the ‘No Control’ Scenario: Potential Damage Costs of Dvv. in Europe ». Journal of Applied Entomology 134 (5): 385–94. https://doi.org/10.1111/j.1439-0418.2010.01510.x.

