## Supplementary material for "PREDATION CAPACITY OF SOIL-DWELLING PREDATORY MITES ON TWO GROUND BEETLE IMMATURE STAGES: A BIOLOGICAL CONTROL PERSPECTIVE FOR TWO MAJOR MAIZE PESTS": Predation tests on WCR broken eggs

1) Discussion: Predation rate for the three species of predatory mites on WCR broken eggs

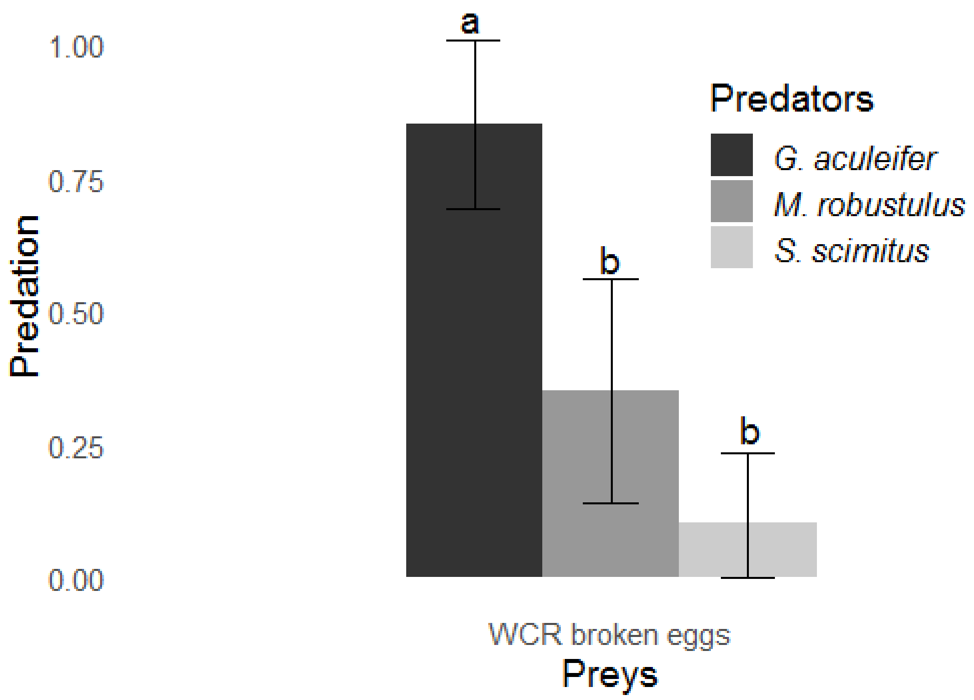
